# Performance comparison of TCR-pMHC prediction tools reveals a strong data dependency

**DOI:** 10.1101/2022.11.24.517666

**Authors:** Lihua Deng, Cedric Ly, Sina Abdollahi, Yu Zhao, Immo Prinz, Stefan Bonn

## Abstract

The interaction of T-cell receptors with peptide-major histocompatibility complex molecules plays a crucial role in adaptive immune responses. Currently there are various models aiming at predicting TCR-pMHC binding, while a standard dataset and procedure to compare the performance of these approaches is still missing. In this work we provide a general method for data collection, preprocessing, splitting and generation of negative examples, as well as comprehensive datasets to compare TCR-pMHC prediction models. We collected, harmonized, and merged all the major publicly available TCR-pMHC binding data and compared the performance of five state-of-the-art deep learning models (TITAN, NetTCR, ERGO, DLpTCR and ImRex) using this data. Our performance evaluation focuses on two scenarios: 1) different splitting methods for generating training and testing data to assess model generalization and 2) different data versions that vary in size and peptide imbalance to assess model robustness. Our results indicate that the five contemporary models do not generalize to peptides that have not been in the training set. We can also show that model performance is strongly dependent on the data balance and size, which indicates a relatively low model robustness. These results suggest that TCR-pMHC binding prediction remains highly challenging and requires further high quality data and novel algorithmic approaches.

## 1. INTRODUCTION

T-cell receptors (TCR) play a crucial role in adaptive immunity mainly through the recognition of peptide fragments from foreign pathogens that are presented by major histocompatibility complex (MHC) molecules. TCRs consist of two transmembrane polypeptide chains, α and β chain; they form a heterodimer on the cell surface. The extraordinary diversity of the TCR repertoire is mainly attributed to a somatic recombination process, V(D)J recombination. Humans can theoretically generate more than 10^15^ different antigen-specific TCRs (1). The diversity of TCR α and β is realized mainly by the complementarity-determining regions (CDRs), with CDR3 being the contact side to the peptide fragment and consequently the most important area for antigen recognition (2) There are two types of MHC molecules, MHC class I and MHC class II molecules, presenting peptides to CD8^+^ and CD4^+^ T cells, respectively.

The major public data resources for TCR-pMHC binding data are VDJdb (3), IEDB (4), McPAS-TCR (5), ImmuneCODE (6), TBAdb (7) and 10X Genomics (8), which all contain TCR CDR3 β chain information.

These are all precious data since identifying cognate TCRs-pMHC binding pairs typically needs both the pMHC multimers technology and single cell sequencing technology (9; 10).

This vast diversity of the TCR repertoire makes it difficult to experimentally cover all possible TCR-pMHC binding pairs. Under the fundamental assumption that the binding between TCR and pMHC is governed by fundamental physicochemical interaction rules, computational approaches could detect and learn those patterns in data. Applying machine learning (ML) and deep learning (DL) approaches to predict the interaction between TCR and pMHC have been explored, resulting in various models such as TITAN, NetTCR, ERGO, DLpTCR and ImRex. Among these models, ERGO and TITAN integrated natural language processing (NLP) techniques, NetTCR-2.0 and ImRex are based on convolutional neural-networks (CNN), and DLpTCR is a combination of CNN, fully connected network (FCN) and deep residual network (ResNet). Unfortunately, to date there exists no appropriate benchmark dataset or workflow to compare contemporary TCR-pMHC prediction models and improve them. In this work, we collected and preprocessed all available major TCR-pMHC data and compared the performance of those state-of-the-art models in different training and testing scenarios.

## 2. RESULTS

### 2.1. Current available data showed a great imbalance

To compare currently available TCR-pMHC prediction models, we first collected data from the most comprehensive public resources, including 10X Genomics, McPAS-TCR, VDJdb, ImmuneCODE, TBAdb and IEDB, then preprocessed separately and afterwards merged into one dataset (TCR preprocessed dataset, tpp dataset). The general process is depicted in Figure 1. The tpp dataset amounts to 113762 entries, out of which 32237 entries contain paired TCR chains, 7167 entries contain only α chains (TRA) and 74358 entries contain only β chains (TRB)(Figure 2A). The composition of the database is shown in Figure 2B. From different data resources, ImmuneCODE contains exclusively TRB information, whereas VDJdb contains the highest number of paired chain examples (Figure 2C). If we further look into the binding pairs between TCRs and peptides presented by MHC molecules, there is a strong imbalance concerning the peptides, i.e. 0.12% of all peptides (20/1659) account for 58.38% of the total entries (66413/113762).

**Figure 1.**
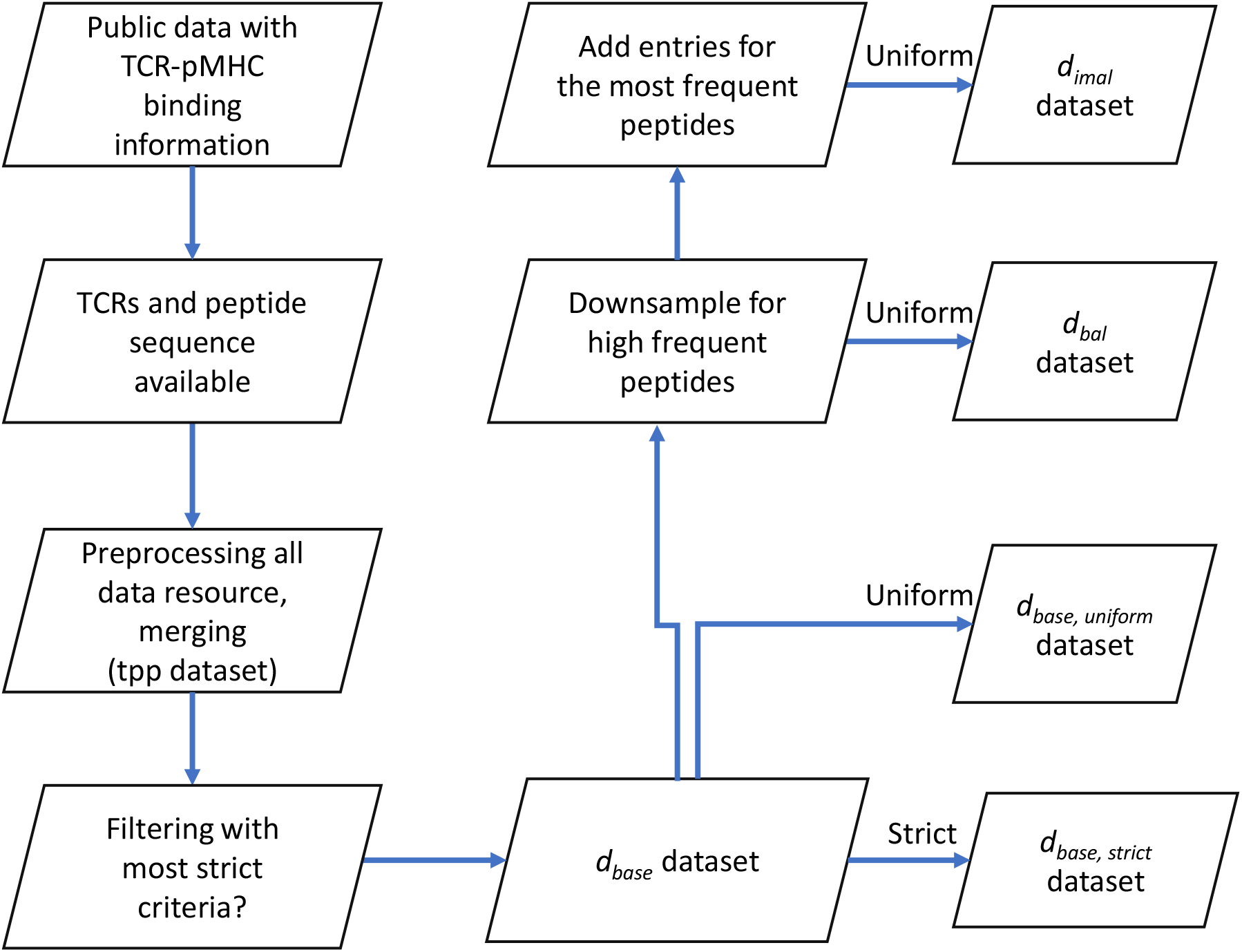
Flow chart shows the basic procedure for preparing different datasets. After collecting data from public resources and merging the preprocessed into one dataset (TCR preprocessed dataset, tpp dataset), different filtering criteria were applied to obtain the positive examples for *d*_*base,strict*_, *d*_*base,uniform*_, *d*_*bal*_ and *d*_*imbal*_ datasets. Negative examples were generated within folds (refer to 4.1.3) after splitting (refer to 4.1.2) to obtain the complete datasets. *d*_*base*_: the base dataset filtered from tpp dataset. *d*_*base,strict*_: strict splitting used on *d*_*base*_. *d*_*base,uniform*_: uniform splitting used on *d*_*base*_. *d*_*bal*_: the balanced dataset filtered from *d*_*base*_, then split using uniform splitting. *d*_*imbal*_: the imbalance dataset filtered from *d*_*base*_, then split using uniform splitting.

**Figure 2.**
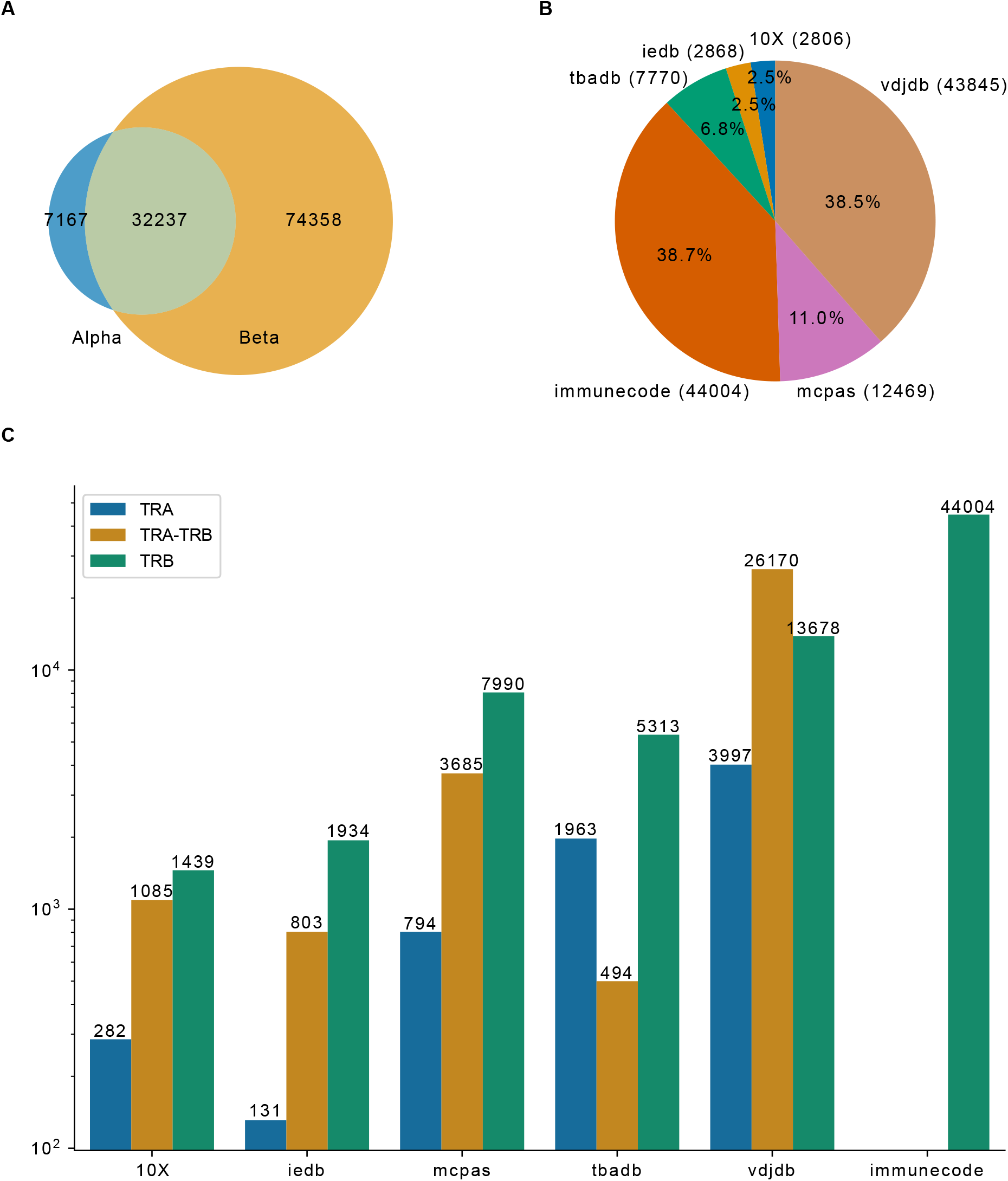
Overview of TCR-pMHC binding data merged from different resources. A) Venn plot shows the overlap of entries that contain only TRA, paired chains or only TRB. The size of the ellipses correlate to the number of entries for each category. B) Pie plot shows the composition of the merged database. Number of entries in each resource indicated in the parentheses. C) TRA and TRB availability for the six major resources.

In order to compare the performance of TITAN, NetTCR-2.0, ERGO, DLpTCR and ImRex, they need to be trained and tested on the same data. We constructed a base dataset (*d*_*base*_), which fulfills all the requirements from these models so that every model can be trained and tested on it. The criteria are: 1) peptide length equals to 9; 2) CDR3 TRB length in the range of 10 to 18; 3) peptides are presented by the HLA-A*02 MHC allele. After applying these criteria, we removed duplicates based on the CDR3 TRB and peptide, this resulted in a total of 15331 entries for *d*_*base*_, across 15039 CDR3 TRB and 691 peptides. The data in *d*_*base*_ is highly imbalanced towards high frequent peptides, 82.66% (12672) of all entries are derived from the top 20 most frequent peptides. The total entries for the top 20 peptides in *d*_*base*_ is shown in Figure 3A. The imbalance of TCRs pairing with the top 20 peptides is highlighted in Figure 3B. The top 20 peptides are paired with 82.66% of the total TCRs while the remaining peptides are paired with the remaining 17.34% TCRs. Furthermore, 517 out of the total 691 peptides have less than five examples per peptide in *d*_*base*_.

**Figure 3.**
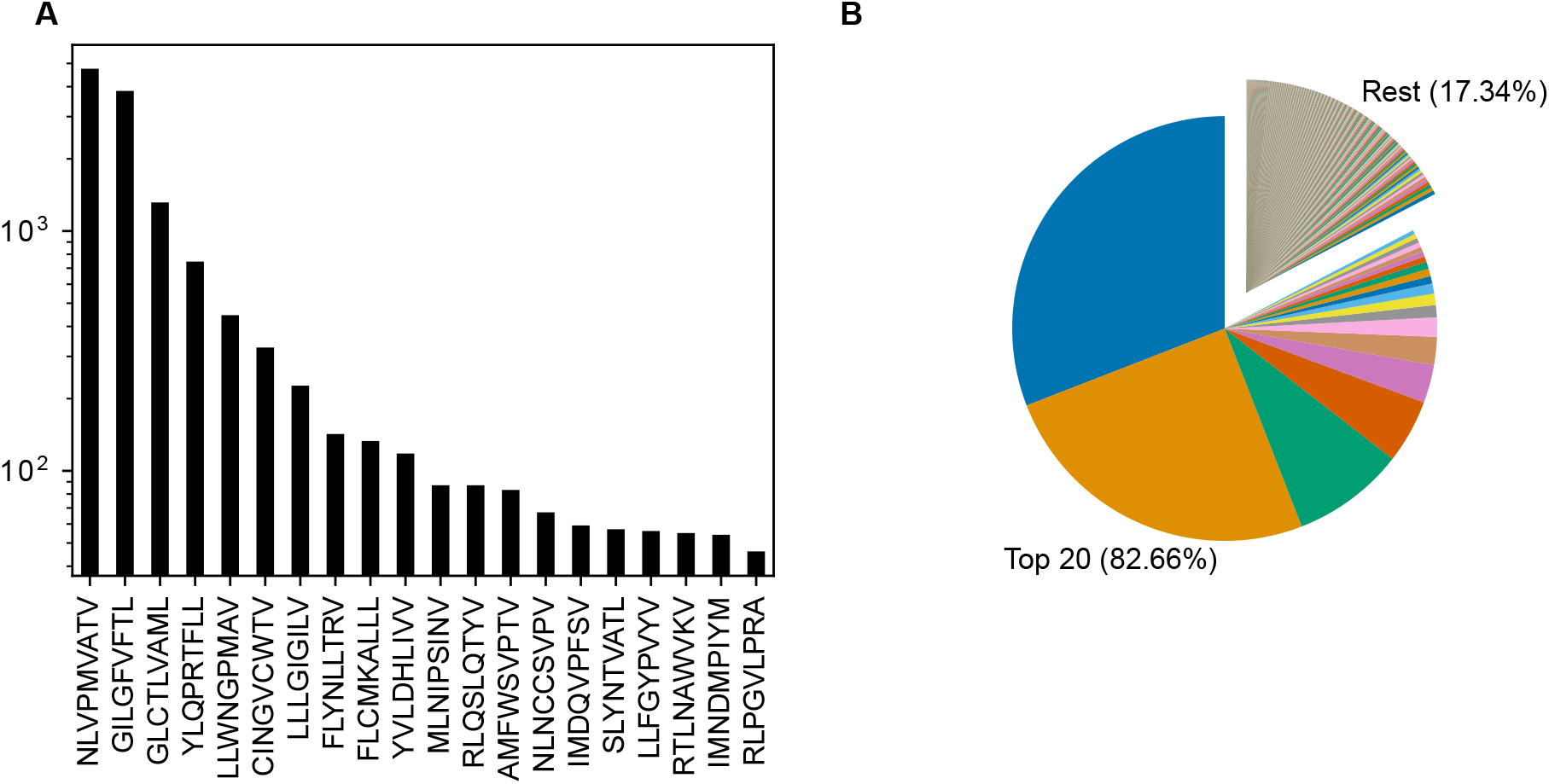
Overview of TCR-pMHC positive binding examples for *d*_*base*_. A) Barplot shows the number of entries for the top 20 peptides. B) Pie chart shows the constitution of examples for the top 20 peptides vs. the rest in *d*_*base*_.

### 2.2 Comparison of model performance on d_base_ indicates that current DL models perform similarly well regardless of model complexity

After acquiring the merged dataset and filtering with the most strict requirements of all tested models we obtained the *d*_*base*_ dataset. In the creation of *d*_*base*_ dataset there were two steps necessary. First, we split the data into five folds as we use 5-fold cross-validation. We used two different splitting methods (see subsection 4.1.2), uniform splitting which keeps the peptide distribution equal across all folds and strict splitting which keeps the peptides in each folds unique. The second prerequisite was to generate negative examples (see subsection 4.1.3), i.e. by assigning combinations of CDR3 β sequences and peptides that do not bind to each other.

Next, we tested six different DL models from five publications. The chosen models predict the binding between a given TCR-pMHC pair. The feature input are the CDR3 TRB sequence of the TCR, and the amino acid (aa) sequence of the peptide. The six models differ in their approaches to embed and process the given features. This subsection compared the different approaches and measured their performance. Models were trained and tested on *d*_*base*_ using 5-fold cross-validation. In Table 1 the tested models are summarized.

**Table 1.**
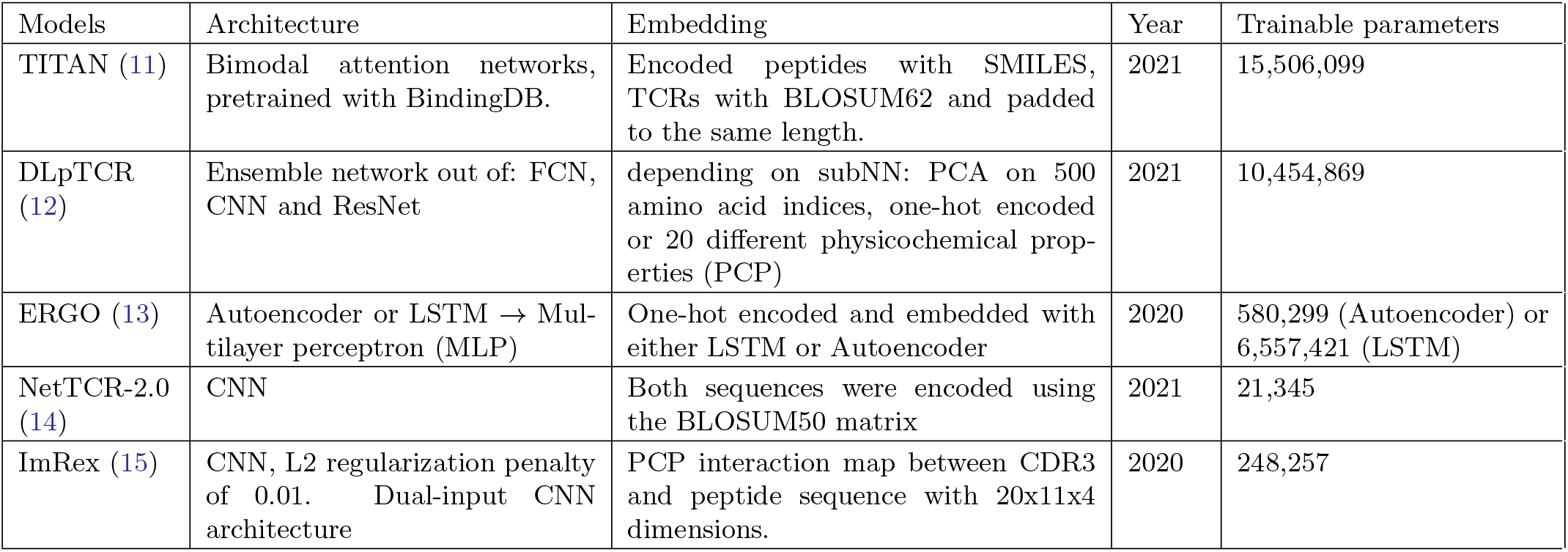
Overview of the tested models.

| Models | Architecture | Embedding | Year | Trainable parameters |
| --- | --- | --- | --- | --- |
| TITAN (11) | Bimodal attention networks, pretrained with BindingDB. | Encoded peptides with SMILES, TCRs with BLOSUM62 and padded to the same length. | 2021 | 15,506,099 |
| DLpTCR (12) | Ensemble network out of: FCN, CNN and ResNet | depending on subNN: PCA on 500 amino acid indices, one-hot encoded or 20 different physicochemical properties (PCP) | 2021 | 10,454,869 |
| ERGO (13) | Autoencoder or LSTM → Multilayer perceptron (MLP) | One-hot encoded and embedded with either LSTM or Autoencoder | 2020 | 580,299 (Autoencoder) or 6,557,421 (LSTM) |
| NetTCR-2.0 (14) | CNN | Both sequences were encoded using the BLOSUM50 matrix | 2021 | 21,345 |
| ImRex (15) | CNN, L2 regularization penalty of 0.01. Dual-input CNN architecture | PCP interaction map between CDR3 and peptide sequence with 20x11x4 dimensions. | 2020 | 248,257 |

The number of trainable parameters (Table 1) of a model indicates the model complexity. We do not see a correlation between the number of trainable parameters and performance of the model. We used a 1:1 ratio of positive:negative binding examples for both training and testing sets. The ROC-AUC score of each model on *d*_*base*_, except for ERGO with the embedding of long short-term memory (LSTM), were above fifty percent (Figure 4). Therefore, almost all models predicted the outcome of a given TCR-pMHC pair better than random guessing. With the exception of ERGO with the LSTM embedding, no ROC-AUC score stood out and performances of those models were within 0.66 *±* .04 ROC-AUC.

**Figure 4.**
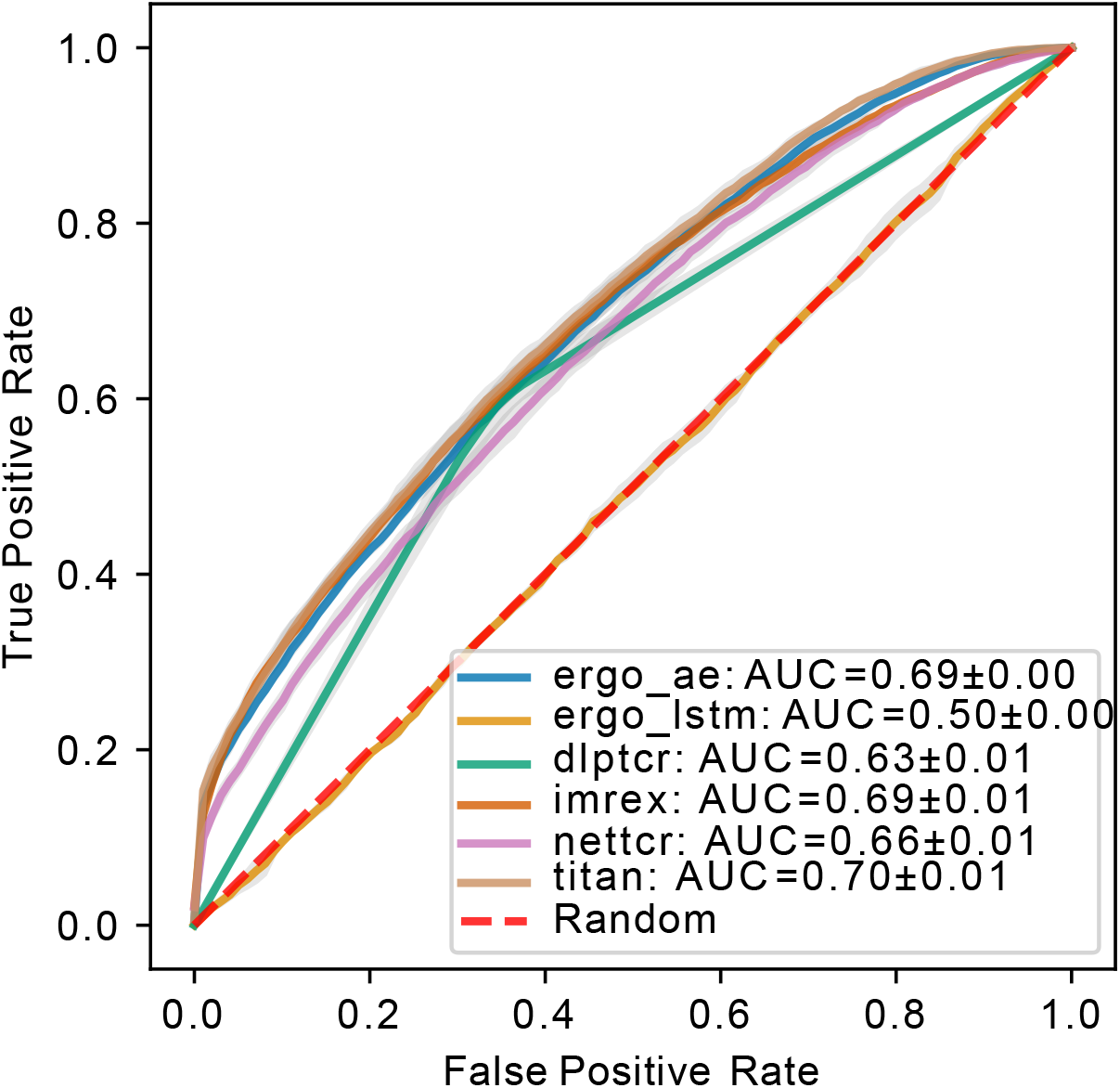
ROC of models predicting binding of TCR-pMHC trained and tested on *d*_*base*_ using uniform splitting. The dashed red line indicates performance of random guessing. ROC curve for DLpTCR looks “linear”, because DLpTCR outputs a binary and not a continuous probability.

### 2.3. Model performance on uniform or strict split data indicates that current models do not perform well on unseen peptides

A generalized prediction model will find interaction patterns that are transferable to new TCR-pMHCs examples. We used two training and testing splitting methods (see subsection 4.1.2) to generate uniform and strict splitting data sets. The main difference of uniform splitting and strict splitting is whether the peptide in the testing set appears in the training set. In uniform splitting the peptides in the testing set also exist in the training set (seen peptides), whereas the peptides in strict splitting have no overlap between training and testing set (unseen peptides). For a generalized TCR-pMHC binding prediction model, it should be able to predict binding on unseen peptides.

The model performance for all models using these two splitting methods is compared in Figure 5. DLpTCR returns a binary in its prediction, and this explains why the curves for DLpTCR in Figure 5 only connect three points. Every other model outputs a value between zero and one, which serves as a probability for the given TCR-pMHC pair to bind. A continuous probability value can generate more points in the ROC and PR curve, if one vary the threshold for a binding and unbinding prediction. Model performance collapsed for strict splitting (comparing Figure 5A and Figure 5C for each model), indicating that current models do not generalize to unseen peptides.

**Figure 5.**
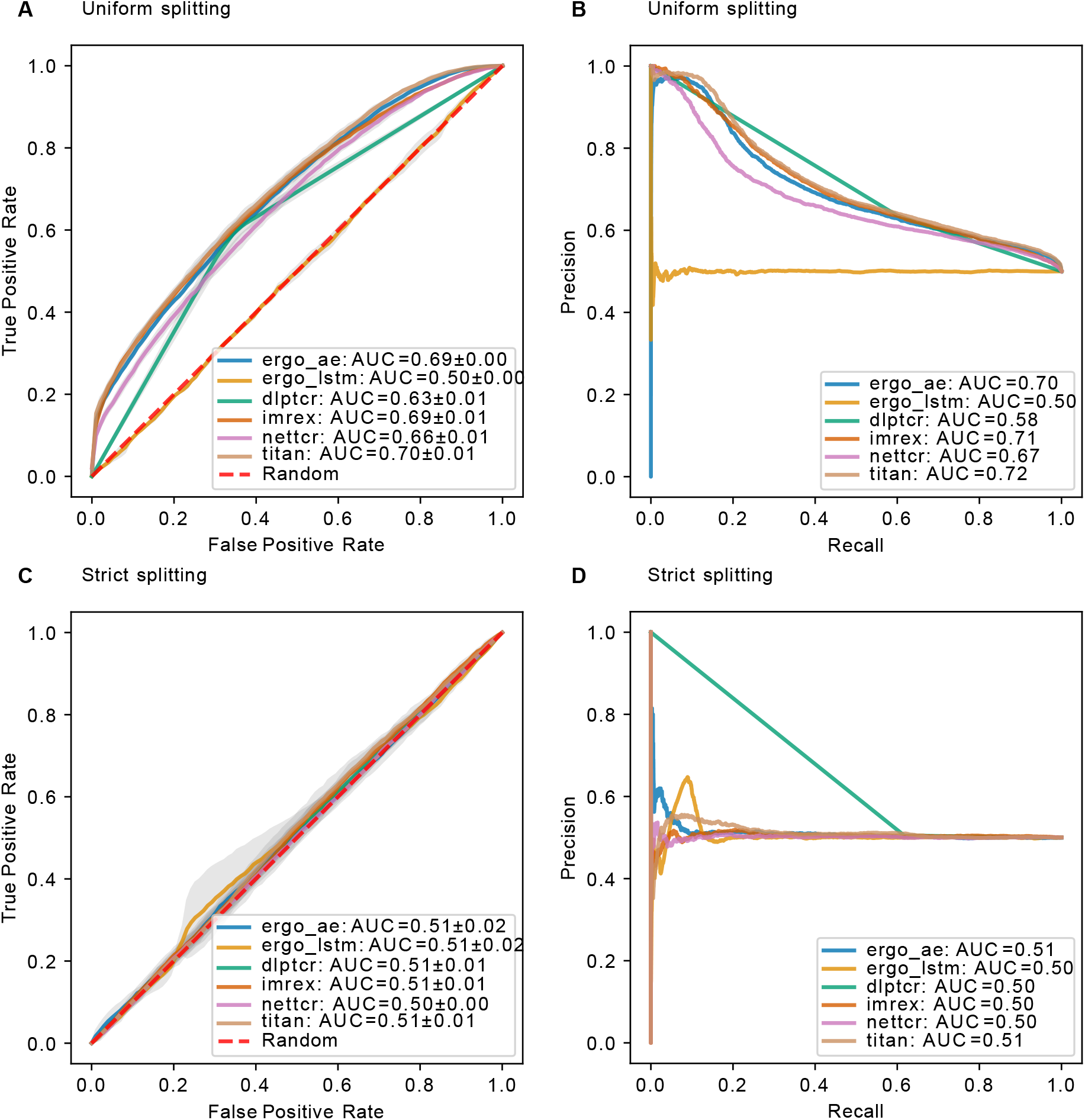
Model performance on datav1 using different splitting methods. A) ROC curve and B) PR curve for models using uniform splitting. C) ROC curve and D) PR curve for models using strict splitting. The dashed red lines indicate performance of random guessing. ROC and PR curve for DLpTCR looks “linear”, because DLpTCR outputs a binary and not a continuous probability.

### 2.4. Collapsing performance on d_bal_ suggests that 5-10 examples per peptide is not sufficient for training state-of-the-art DL models

After comparing the results for *d*_*base*_ using uniform/strict splitting, we realized that current models are not able to predict the binding for unseen peptides. Since results for uniform splitting showed moderate prediction ability, we suspected that these models learned for the high frequent peptides. In order to elucidate this, we prepared a new balanced data set (*d*_*bal*_) to test this hypothesis. Based on *d*_*base*_, we filtered out entries with less than 5 examples per peptide and afterwards we downsampled 4.1.4 each unique peptide, so that each peptide in *d*_*bal*_ only contains 5-10 examples. This resulted in *d*_*bal*_ with a total of 2812 examples, across 1397 unique CDR3 TRB sequences and 174 unique peptides. Training the models on *d*_*bal*_, we saw a complete collapse of performance for the models (Figure 6), similar to *d*_*base*_ strict splitting. This indicates either that 5-10 examples per peptide is not sufficient for a predictive model to learn the general TCRs-pMHC binding rules or that a total of 2812 examples is not enough to train and test the models on. In the following subsection we investigated how data imbalance impacted the model performance.

**Figure 6.**
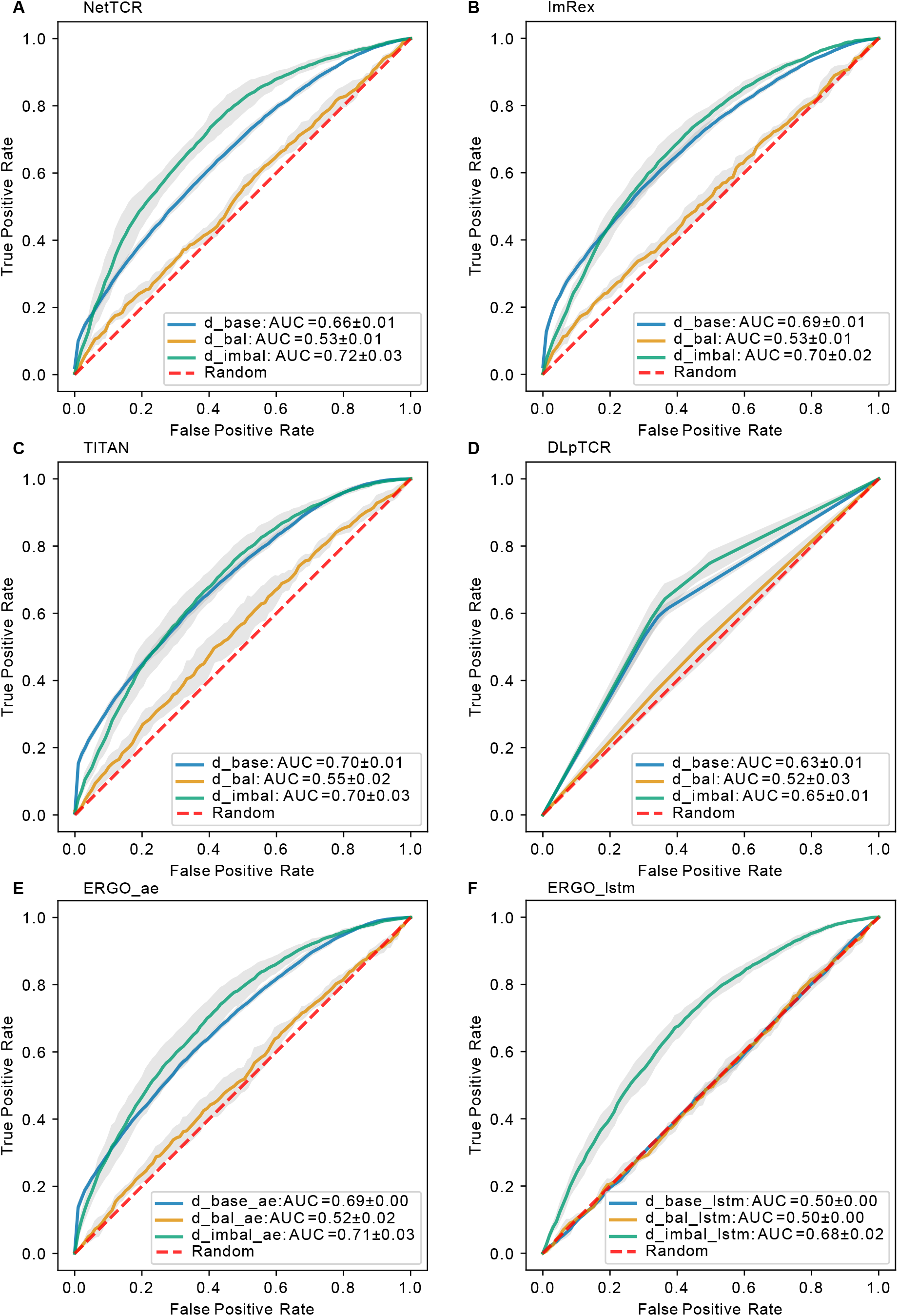
Model performance on different datasets using uniform splitting. ROC curve for A) NetTCR-2.0, B) ImRex, C) TITAN, D) DLpTCR (Curves looks “linear”, because DLpTCR outputs a binary and not a continuous probability), E) ERGO autoencoder model and F) ERGO LSTM model using *d*_*base*_, *d*_*bal*_ and *d*_*imbal*_. The dashed red diagonal line indicates performance for random guessing.

### 2.5. Model performance comparison on d_base_ and d_imbal_ indicates that “success” is only due to the most frequent peptide

The difference between *d*_*bal*_ and *d*_*base*_ is in the size and the imbalance regarding the peptide distribution. The degree of balance can be calculated with the formula for Shannon entropy,

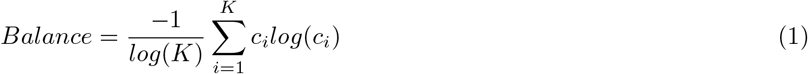

with *K* as the number of unique peptides and *c*_*i*_ as the occurrence in percentage for peptide *i*. We constructed *d*_*imbal*_ to investigate whether the peptide imbalance or the data size impacts the performance more. This dataset included all available data for the most frequent peptide (mfp) (“NLVPMVATV”), but filtered and downsampled the remaining peptides (non-mfp). In total, *d*_*imbal*_ has 12268 entries, with 7678 unique CDR3 TRB sequences and 174 unique peptides. This dataset has a higher peptide imbalance than *d*_*base*_ and a smaller size (see Table **??**).

We would expect *d*_*base*_ which contains more input data to have a better performance over *d*_*imbal*_ if the model can learn a general binding rule. However, models trained on *d*_*imbal*_ had a prediction power comparable to models trained on *d*_*base*_, and even slightly better than models trained on *d*_*base*_ (Figure 6). In the case of ERGO with LSTM embedding, which was as bad as random guessing if trained on *d*_*base*_, if trained on *d*_*imbal*_ we saw an increase in prediction performance. Therefore, we conclude that peptide imbalance impacts the performance more than the size of the data. This result also suggests that all models learned the binding rule for the most frequent peptide examples.

### 2.6. Performance increases with peptide imbalance

Next, we investigated whether the learned most frequent peptides from *d*_*imbal*_ can be transferred to predict the binding for less frequent peptides. Overall, the ROC-AUC scores for the models trained on *d*_*imbal*_ were significantly higher than the one trained on *d*_*bal*_ (Figure 6). If models trained on *d*_*imbal*_ also showed better performance for the non-mfp, compared to models trained on *d*_*bal*_, this would mean that the learned mfp increases the likelihood of generalization. In Figure 7, we compared the accuracy on non-mfp data using models trained on the two datasets and no change in performance was observed. We observed a strong data dependency regarding the performance of all models. In retrospect, the success of previously published models could thus be attributed to the peptide imbalance within each dataset.

**Figure 7.**
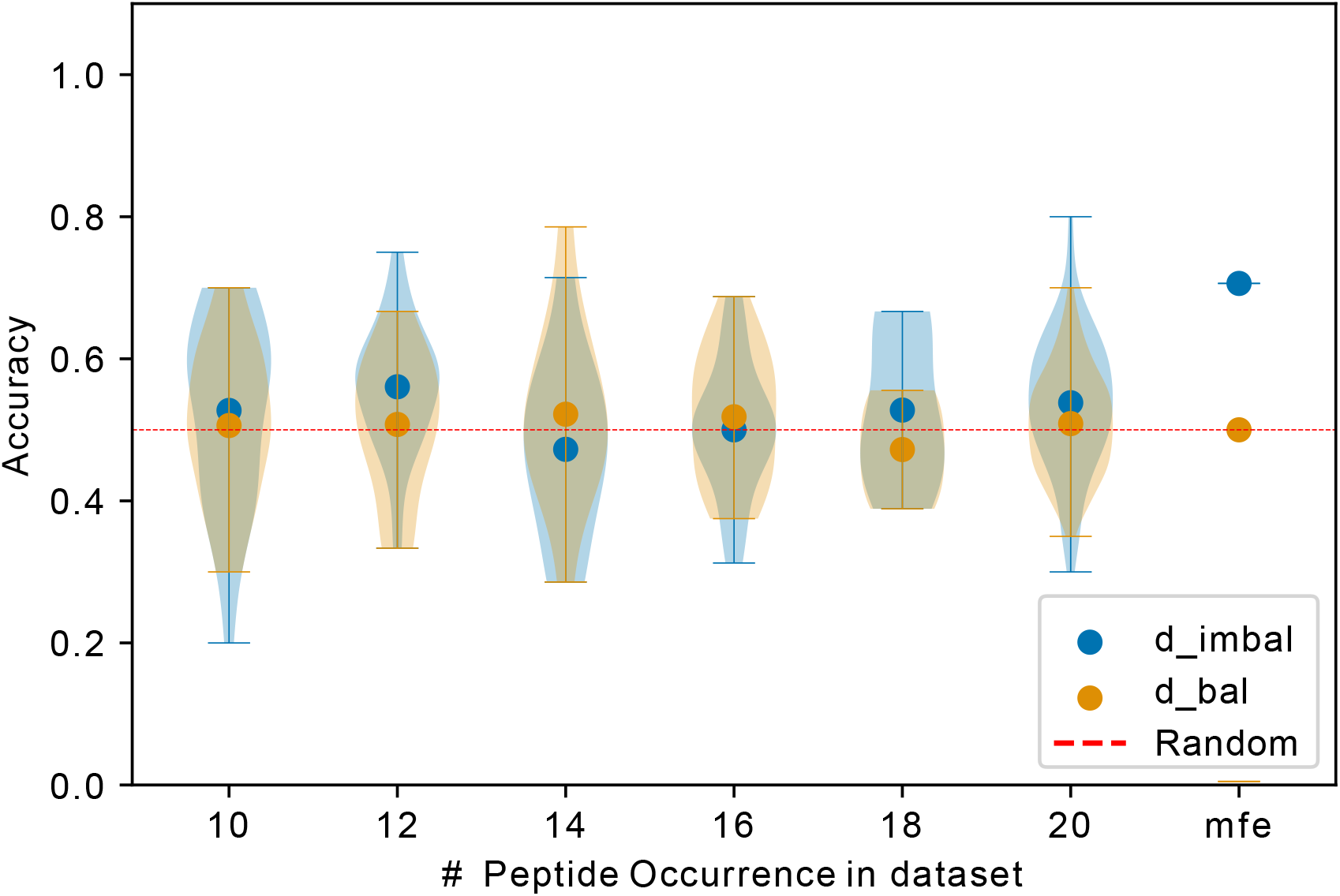
Exemplary comparison of NetTCR-2.0 performance trained on *d*_*imbal*_ and *d*_*bal*_. Data points indicate accuracy for models (trained on different datasets) testing on unique peptide with different occurrence. mfp: most frequent peptide 20 examples in *d*_*bal*_ and 9476 examples in *d*_*imbal*_.

## 3. DISCUSSION

In this work, we compared different state-of-the-art models for the prediction of TCR-pMHC binding. By using different train/test splitting methods, we were able to contrast the performance of the models between unseen and seen peptides. Our findings clearly show that all models with different complexity fail to predict on unseen peptide examples. This is consistent with the findings of Grazioli et al. (16), who contrasted the performance between uniform and strict splitting as well. They show that ERGOII as well as NetTCR2.0 performs worse in strict splitting. Here, we have also tested NetTCR2.0 and a predecessor model of ERGOII (ERGO), but additionally includes TITAN, DLpTCR and ImRex to cover all the current state-of-the-art models (Table 1) for TCR-pMHC binding prediction. We showed that the performance stays the same across models with different complexity. Notably, Grazioli et al. suggested that TITAN is a potential candidate to have a generalized prediction prowess. TITAN (11) by Weber et al. applied strict splitting themselves and measured a performance of up to 0.62 ROC-AUC. However, we could not replicate this result based on our dataset. A possible explanation why Weber et al. measured a better performances could be that they only used data from VDJdb (peptides from various origin) and ImmuneCODE (exclusively COVID data). Merging those two datasets will result in mostly peptides associated with COVID (105/192 [54.69%] assuming VDJdb does not contain many COVID data). Even if the peptides in the testing and training sets are disjoint in strict splitting, there might be similar peptides across the training and testing set, due to their same origin from COVID. This may have contributed to the better performance reported. If this hypothesis is true, given enough training examples, it might be possible for TITAN and other models to not only predict peptide-specific binding but also origin-specific binding.

We also investigate the impact of peptide imbalance on the performance of the models. To the best of our knowledge, we have not seen similar training and testing of the models on different data scenarios (*d*_*base*_, *d*_*bal*_ and *d*_*imbal*_). The data scenarios vary in size and peptide distribution. We suggest that peptide imbalance contributes more to a better performance of the models than size.

This is consistent with the consensus that currently available data are not sufficient, an issue raised so far by every study of these models (12; 14; 13; 15; 11). Our results support the idea that a generalized predictive model requires data that is not only large but also massively diverse to uncover a large range of potential pMHC-TCR binding rules.

The hypothesis that models such as TITAN might be able to predict unseen but similar peptides or peptides from the same origin is a very interesting research question for future work. If this hypothesis holds, we need a global effort to experimentally screen a set of peptides to cover a diverse peptides pool, and make use of the generated data for constructing a generalizable prediction model.

A limitation of this study is that our datasets only comprised TCRs from CD8^+^ T cells pairing with peptides presented by the HLA-A*02 allele without considering other MHC alleles, however, it was important to exclude additional variables such as HLA isotypes at this point. Moreover, we only compared DL models for predicting binding between random TCRs and random pMHC, not epitope-specific models (i.e. prediction whether random TCRs bind to a specific peptide). Meysman et al. have compared superficially different approaches to TCR-pMHC binding (17), but also raised the importance of a truly independent benchmark. They reveal that additional information like CDR1/2 improved the prediction, but they did not investigate the role that imbalance, size or overtraining might have on model performance by using those additional features within the used dataset.

## 4. METHODS

### 4.1. Data preprocessing

#### 4.1.1. Date merging and preprocessing

We downloaded the data from six different resources. We unified the column names of (CDR3 TRA, CDR3 TRB, peptide and MHC, etc.). We only kept entries that have a peptide and at least either a CDR3 TRA or TRB sequence. Only TCRs sequences and peptide sequences that use the 20 valid amino acid residues are kept. After this quality control, all data from different resources were merged into one dataset (tpp dataset), duplicates in this merged dataset were then removed. The preprocessing of the merged dataset and pre-filtering for different datasets are shown in Figure 1.

#### 4.1.2. Splitting

We explored two different splitting methods (Figure **??**). The first method kept the distribution of the peptide in each part (uniform splitting). The second method distributed peptides to each part, so that no peptide is in two different parts (strict splitting). The strict splitting we used here is inspired by the splitting method from the TITAN (11) model. Strict splitting was only used for *d*_*base*_ (Figure 1). *d*_*base,strict*_ and *d*_*base,uniform*_ vary in size (Table **??**), because strict splitting includes peptides with less than five examples. In subsection 2.1 we showed a data imbalance in peptides. For the 5-fold cross-validation in strict splitting we ensured, that each fold won’t have a peptide exceeding more than half of it’s entries. If a peptide has more entries it will be downsampled to the half of the fold size. Uniform splitting exclude peptides with less than five examples, because uniform splitting requires at least one example for each peptide in all five folds. Table **??** shows that *d*_*base,strict*_ have more unique peptides but less total entries compare to *d*_*base,uniform*_. In *d*_*base,strict*_, we downsampled many positive examples (for high frequent peptides) in order to generate negative examples within each fold without external reference TCR repertoire, this reduces the total number of examples in the dataset, while in *d*_*base,uniform*_, some examples for less frequent peptides were filtered out to ensure at least one example in each fold.

#### 4.1.3. Negative example generation

The negative example data were created by rearranging non-matching TCR-pMHC pairs. Each peptide got a new CDR3 TRB partner, which originally binds to other peptides. Alternatively, we used additional reference TCR repertoire, to pair with the pMHC in order to get negative TCR-pMHC binding examples for *data*_*imbal*_.

#### 4.1.4. Downsampling

Peptides are not uniformly distributed throughout tpp dataset. Some peptides occur only a few times (low frequent peptides) and some occur hundreds of times (high frequent peptide). For *d*_*bal*_ and *d*_*imbal*_ we downsampled the high frequent peptides to keep only 10 random examples for each peptide.

### 4.2. Model performance measurement

We downloaded the source code for all models from their respected GitHub repository. We evaluated all models with 5-fold cross-validation. We used our datasets to train the models with the default parameters. The performance is measured by the area under the receiver and operator curve (ROC-AUC) (18), as well as the area under the precision recall curve (PR-AUC)(19). The best ROC-AUC models was saved and evaluated on testing set.

## 5. DATA AVAILABILITY

All data used in this manuscript can be found at https://github.com/imsb-uke/TCRs-pMHC. The 10X Genomics dataset can be found at https://www.10xgenomics.com/resources/application-notes/a-new-way-of-exploringimmunity-linking-highly-multiplexed-antigen-recognition-to-immune-repertoire-andphenotype/. The McPAS-TCR dataset can be found at http://friedmanlab.weizmann.ac.il/McPAS-TCR/, the VDJdb dataset can be found at https://vdjdb.cdr3.net/, the ImmuneCODE dataset can be found at https://clients.adaptivebiotech.com/pub/covid-2020, the TBAdb dataset can be found at https://db.cngb.org/pird/tbadb/ and the IEDB dataset can be found at http://www.iedb.org/. Final merged data can be found at https://github.com/imsb-uke/TCRs-pMHC.

## 6. ACKNOWLEDGEMENTS

This work was funded by grants from the Deutsche Forschungsgemeinschaft (DFG), grants CRC1192, project number 264599542 and PR727/14-1, project number 497674564.

