## supplemental file for "Performance comparison of TCR-pMHC prediction tools reveals a strong data dependency"

### 1. SUPPLEMENTARY INFORMATION

**Table S1.** Dataset overview

| | $d_{base,strict}$ | $d_{base,uniform}$ | $d_{bal}$ | $d_{imbal}$ |
|---|---|---|---|---|
| entries | 15964 | 28716 | 2812 | 12268 |
| unique peptide | 691 | 174 | 174 | 174 |
| unique CDR3 Beta | 7805 | 14141 | 1397 | 7678 |
| Shannon entropy | 0.65 | 0.49 | 0.99 | 0.33 |

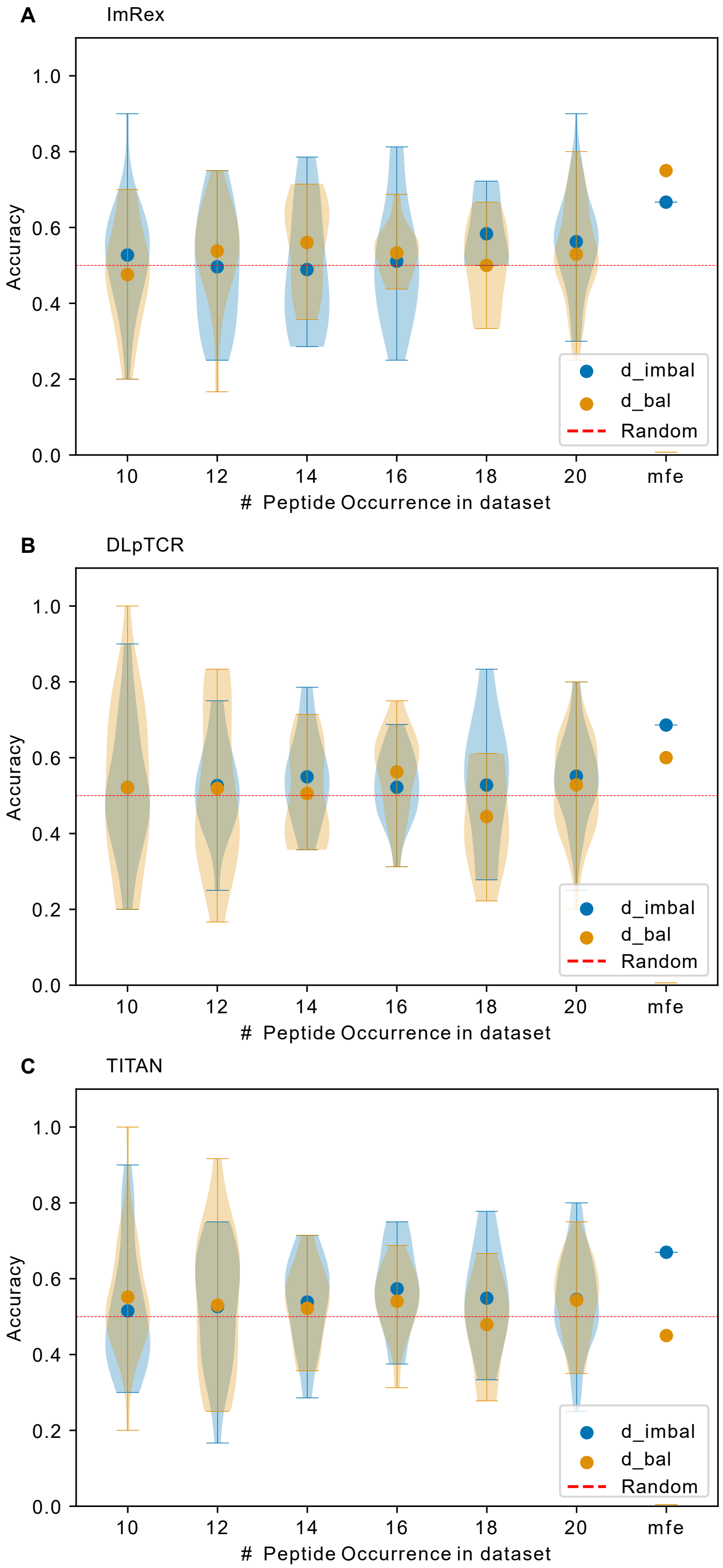


**Figure S1.** Performance trained on $d_{imbal}$ and $d_{bal}$ for A) ImRex, B) DLpTCR and C) TITAN. Data points indicate accuracy for models (trained on different datasets) testing on unique peptide with different occurrence. mfe: most frequent peptide 20 examples in $d_{bal}$ and 9476 examples in $d_{imbal}$.

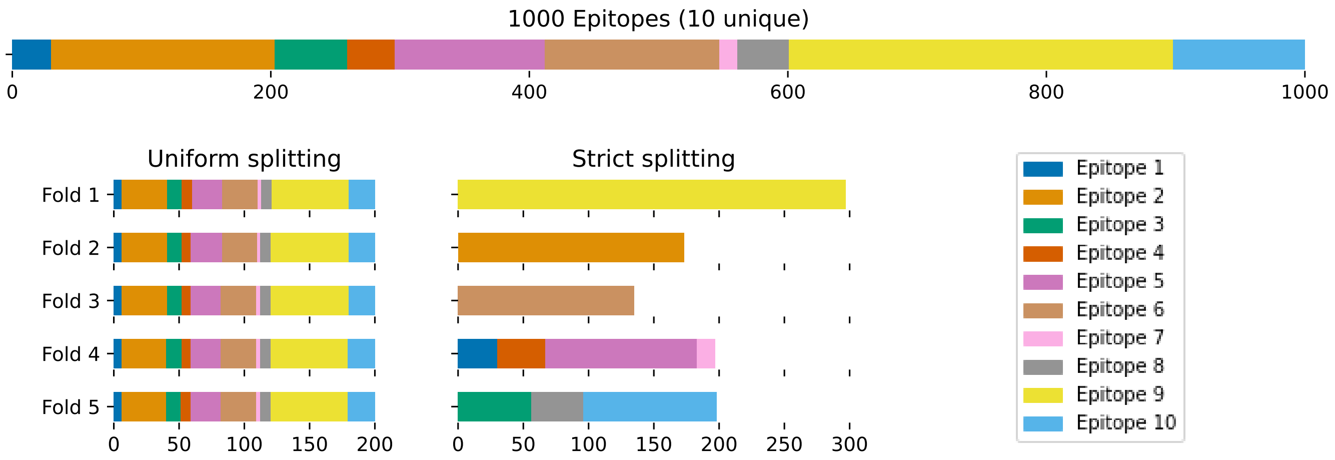


**Figure S2.** Uniform and strict splitting schematically demonstrated with an imbalanced dataset of 1000 entries and 10 unique peptides.
